# FoundedPBI: Using Genomic Foundation Models to predict Phage-Bacterium Interactions

**DOI:** 10.64898/2026.03.24.713871

**Authors:** Pere Carrillo Barrera, Arthur Babey, Carlos Andrés Peña

## Abstract

The scalability of phage therapy as a viable alternative or complement to antibiotics is limited by the labor-intensive experimental screening required to identify compatible phage-bacterium pairs. To accelerate this discovery process, we propose FoundedPBI, an ensemble deep learning approach that leverages the emergent capabilities of genomic foundation models, large language models pre-trained on vast DNA corpuses to predict phage-bacterium interactions from DNA sequences alone. We employ an ensemble strategy that aggregates outputs from three state-of-the-art DNA language models into a unified meta-embedding, which is then processed by a neural classifier. Our approach makes two key contributions: (1) We demonstrate that performing ensemble learning across models trained on different genomic data—i.e., prokaryotic (Nucleotide Transformer v2, DNABERT-2) and bacteriophage (MegaDNA) genomes—captures partially-orthogonal biological signals, yielding 6% F1-score improvement over the best individual model. (2) We adapt long-context NLP aggregation strategies to handle whole bacterial and phage genomes (up to 5M base pairs) that exceed the foundation models’ context windows (12-96K bp) by a factor of 50–100, a critical challenge largely unaddressed in prior genomic deep learning work. On the PredPHI benchmark, FoundedPBI achieves a 76% F1-score outperforming the current state-of-the-art (PBIP) by 7%. On our internal dataset (CI4CB), we achieve 93% F1-score, improving our previous best methods by 4%. These results demonstrate that ensemble learning with proper long-context handling enables effective knowledge transfer of genomic foundation models to specialized prediction tasks.

## Introduction

Antimicrobial resistant, or multi-drug resistant (MDR), bacteria are quickly becoming a major threat to humanity. Studies predict that they will cause 40 million annual fatalities in the coming years (Oyenuga et al., 2024). Having developed resistance to the most common antibiotics, these bacteria are very difficult to eliminate and are growing stronger. Recently, phage therapy has regained considerable attention as a complementary approach to antibiotics, as it uses highly specific viruses-phages-that can target and combat MDR bacteria with almost no undesirable side effects (Barron, 2022). However, finding adequate phages that infect a specific target bacterium is a difficult task, due to the huge amount of distinct phages available and their narrow host range. A common hypothesis in the field, generally accepted as true since the last century (Luria and Delbrück, 1943), is that enough information about their possible interaction is contained in their genomes. So, predicting interactions only from bacterial and phage DNA sequences has become a common approach (Ma et al., 2025; Zhou et al., 2024a). It is in this context that the Phage-Bacterium Interaction (PBI) problem is defined.

A common recent approach for PBI prediction is to delegate feature extraction to embedding models that create vector representations of the sequences. For instance, Zhou et al. (2024a) use the Hyena architecture to extract embeddings from viral sequences. They then apply contrastive learning to map these embeddings into the same space as prokaryotic ones, grouping related organisms together and pushing unrelated ones apart. Ma et al. (2025) go a step further by using a pre-trained protein embedding model based on LSTM called UniRep (Alley et al., 2019), to create representations for both bacteria and bacteriophages, on which the final prediction is based.

In this paper, we present FoundedPBI, a new approach to PBI modelling. It uses existing general-purpose DNA foundation models to create embeddings for bacterial and bacteriophage genomes and classifies each pair using a neural network, attaining better results on benchmark datasets than the current best models. FoundedPBI shows that general-purpose models can address the PBI problem with minimal resources, eliminating the need for fine-tuning or adaptation. It also shows that knowledge from multiple foundation models can be combined to create more precise embedding representations, allowing for a modular approach to FoundedPBI architecture.

### Ensemble Learning and Long-Context Handling

While genomic foundation models have demonstrated strong performance in several DNA sequence analysis tasks, their application to PBI prediction faces two fundamental challenges that are not currently well addressed.

Firstly, existing foundation models differ substantially in terms of their training data, architectures, and inductive biases. Some are trained on extensive genome collections spanning multiple species that explicitly exclude viral sequences, while others are trained exclusively on bacteriophage DNA. These models also employ fundamentally different architectures each imposing distinct inductive biases on the learned representations. We hypothesise that these differences result in complementary embeddings that capture different aspects of phage and bacterial biology. Rather than selecting a single “best” model, we propose an ensemble learning approach that combines multiple perspectives through meta-embedding concatenation (Dietterich, 2000), allowing the downstream classifier to leverage a wide spectrum of genomic features.

Secondly, a critical challenge is a severe length mismatch between genomic sequences and model context windows. While bacterial genomes average 5 million base pairs (bp) and phage genomes range from 50 to 200 Kbp, current genomic foundation models only support context windows of 12–96 Kbp. This means that a model can only observe a very small part of a complete genome in a single forward pass (0.1–10% for bacteria). To address this, we adopt embedding strategies from NLP (Sannigrahi et al., 2023), which face similar challenges when embedding long documents using sentence-level models. We systematically evaluate multiple aggregation approaches, including truncation, averaging, and weighted windowing methods, to identify optimal techniques for preserving genomic information across entire sequences. To our knowledge, this is the first work to rigorously address the long-context problem for genomic foundation models in a biological prediction task.

## Materials and Methods

### Datasets

We evaluate FoundedPBI on two complementary benchmarks. A controlled, proprietary dataset for high-confidence validation and comparison with our previous methods, and an external benchmark to assess generalization and state-of-the-art comparison. In Section 3, we report performance on the following datasets:

- **CI4CB**: Our internal dataset. It enables direct comparison with previous and ongoing modelling efforts (i.e., PERPHECT, Distilled DNABERT), demonstrating iterative improvement in our research program.
- **PredPHI**: (Li et al., 2021) Independent benchmark for comparing against published state-of-the-art methods and evaluating generalization across diverse organisms, serving as our primary validation of the proposed contributions.

#### CI4CB Data

The combination of two different datasets, called herein *public* and *private* and both compiled by Leite et al. (2018), is used for training, validating and testing the models. The *public* dataset was created from public databases, such as GenBank (Sayers et al., 2020) and PhagesDB (Russell and Hatfull, 2016), while the *private* dataset was constructed through controlled research investigating interactions between selected bacteria and phages.

In total, these two datasets contain information about 7721 interactions between 231 bacteria and 3539 bacteriophages. The combination of datasets is split into train and test, containing 80% and 20% of the data, and 10-fold cross validation is used during training. A special effort is made to train and test the three models on very similar splits of the dataset, in order to provide the best comparison possible.

The *private* dataset is significantly unbalanced with almost a 1:6 ratio of positive versus negative interactions. This is expected in experimental phage–host interaction data, where confirmed infections are inherently rarer than non-infections. In contrast, the *public* dataset is built imbalance. To mitigate the skew in the private set, we oversampled the positive class by duplicating instances until a balance was reached. This resulted in a total of 8464 examples for training plus validation, and 2030 instances for testing, equally divided between positive and negative interactions. Oversampling was performed after the train/test split to prevent data leakage.

#### PredPHI Data

To evaluate generalization and compare against published methods, we use the PredPHI benchmark (Li et al., 2021) provided by PBIP (Ma et al., 2025) as *species-level interaction dataset*. It includes 6938 interactions, from 301 bacteria and 3449 phages, which where extracted from the PhagesDB (Russell and Hatfull, 2016) and NCBI GenBank (Sayers et al., 2020) in 2019. They also provide the train and test splits used, which are divided by the submission date of the interactions. Following the protocol established by PredPHI and adopted by PBIP, interactions submitted before 2016 form the test set and those from 2016 onward form the training set. This leaves 5702 interactions for training and 1236 for testing. In both cases, the number of positive and negative instances is balanced.

### Framework Overview

The proposed framework, as seen in Figure 1, is composed of two major blocks: A *non-trainable backbone* and a *trainable classification head*. It is based on the architecture of the *predictor* module of the *PERPHECT* system (Ataee et al., 2020), replacing the convolutional feature extractor with pretrained genomic foundation models, and expanding the classification head.

**Figure 1.**
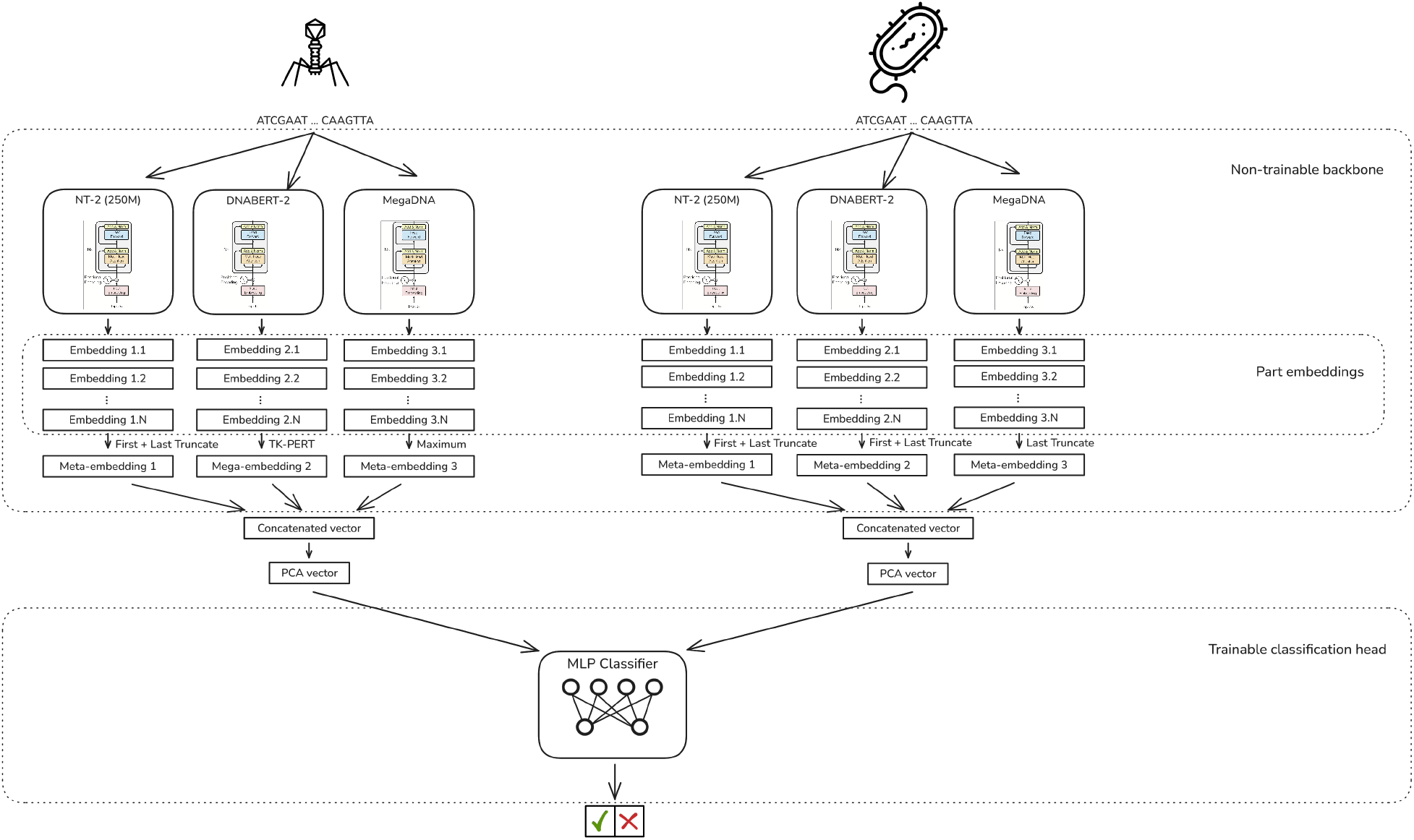
Overview of the proposed framework. It takes advantage of multiple genomic foundation models to create embeddings for the sequences, which are merged and compressed into one meta-embedding from which a classification head makes the final prediction.

### Ensemble Learning and Genomic Foundation Models

Rather than selecting a single genomic foundation model, we construct meta-embeddings by concatenating representations from three architecturally and data-diverse models, as shown in Figure 1. The ensemble is designed so that each model contributes a complementary perspective: different training corpora, architectural inductive biases, and tokenization strategies result in partially independent errors, allowing the ensemble to correct individual model mistakes through implicit majority voting in the embedding space. This hypothesis is validated in Section 3, where we demonstrate that the ensemble consistently outperforms any individual model.

Three state-of-the-art models are used for the current version of FoundedPBI. They were selected to maximize diversity along three axes: training data composition, architecture, and tokenization. Other models such as EVO1 (Nguyen et al., 2024), EVO2 (Brixi et al., 2025), and AIDO.DNA (Ellington et al., 2025) were also considered but discarded due to lack of architectural novelty relative to the selected models or prohibitive resource constraints, as for the EVO family.

#### Nucleotide Transformer v2

The Nucleotide Transformer v2 250M (Dalla-Torre et al., 2024) is an encoder-only transformer designed for genomics data. It tokenizes input sequences into overlapping 6-mers and uses rotary embeddings for positional encoding, with a maximum context length of 12K bp. It was trained on 3 202 human genomes plus 850 genomes from diverse domains (mammals, fungi, bacteria), explicitly excluding viral DNA.

#### DNABERT-2

DNABERT-2 (Zhou et al., 2024b) is the second iteration of the DNABERT model (Ji et al., 2021), based on the BERT architecture. It introduces Byte Pair Encoding (BPE) for tokenization and Attention with Linear Biases (ALiBi) for positional encoding, technically eliminating any hard maximum sequence length. However, because memory consumption grows quadratically with sequence length, the practical input length in our setup is limited to 32K bp. Like Nucleotide Transformer v2, it was trained on genomes from all domains of life except viruses.

#### MegaDNA

MegaDNA (Shao and Yan, 2024) is a generative model based on a GPT-style multiscale three-layer decoder-only transformer (Yu et al., 2023), designed for sequence generation. Following the authors’ recommendations, we extract embeddings by concatenating representations from all three decoder layers, yielding a final embedding of dimension 964. A key advantage over the other two models is its practical maximum context length of 96K bp, allowing substantially more genomic context per embedding. Crucially, MegaDNA was trained exclusively on 100 000 bacteriophage genomes, making it complementary to the host-genome-trained Nucleotide Transformer v2 and DNABERT-2.

### Non-Trainable Backbone

This block is composed of two architecturally equal branches, one responsible for creating a meta-embedding for the bacteriophage DNA sequence and the other for the bacterium sequence.

Each branch uses three different genomic foundation models to create three embeddings for the sequence, which are then joined by concatenation. To embed the long DNA sequences, of up to 10M bp for some bacteria, with these length-limited foundation models, the strategies presented by Sannigrahi et al. (2023) for document embeddings based on sentence embedding models are extended for the genomics language.

### Long-Context DNA Sequence Handling

Embedding the entire DNA sequences of bacteria or phages with the presented foundation models is an impossible task, as almost all sequences are significantly longer than the maximum length allowed for the models. The average sequence length in the used data is 5M bp, more than 50 times longer than the maximum size. This problem has already been explored in the NLP field, when creating embeddings for entire documents. The approach used by FoundedPBI, presented by Sannigrahi et al. (2023) is to use sentence embedding models to embed the document by chunks, and combine them using different strategies.

Following their work, we implemented and tested the best-performing techniques they reported, as their results cannot be used directly since their study was conducted on natural language data. All the techniques embed the DNA sequences into fixed-length chunks and combine them using one of the following strategies.

- **Truncate Strategy:** Only the embedding from the first chunk is used.
- **Bottom Truncate Strategy:** Only the embedding from the last chunk is used.
- **Top + Bottom Truncate Strategy:** The embeddings from the first and last chunks are concatenated to form the resulting meta-embedding.
- **Average Strategy:** All the chunk embeddings are averaged with uniform weights.
- **Maximum Strategy:** The maximum value at each dimension of all the embeddings is used to create the final meta-embedding.
- **TF**_**2**_**/IDF**_**4**_ **Strategy:** All the chunk embeddings are averaged using a variant of TF/IDF weights. To define the concept of “word” in the genomic language, each 6-mer is considered as a different word, following the tokenization technique from Nucleotide Transformer v2.
- **TF**_**4**_**/IDF**_**4**_ **Strategy:** All the chunk embeddings are averaged using another variant of TF/IDF weights.
- **TK-Pert Strategy:** This technique creates multiple smoothed overlapping windows that weight the contribution of each chunk in a sequence according to a modified PERT function, with parameters *J* = 16 (number of windows), *gamma* = 20 following the work of Sannigrahi et al. (2023). The resulting meta-embedding is formed by concatenating all the windows.

We systematically evaluated multiple long-context aggregation strategies via grid search over 1,296 configurations on the CI4CB validation set. Analysis of top-performing combinations (see Supplementary Figure S1) guided our selections: Nucleotide Transformer v2 uses First+Last Truncate for both bacterium and phage branches; DNABERT-2 uses First+Last Truncate (bacterium) and TK-PERT (phage); MegaDNA uses Last Truncate (bacterium) and Maximum (phage). The prevalence of truncation-based strategies suggests that these models capture sufficient taxonomic signal from short genomic fragments, without requiring full-genome context. This is consistent with the foundation models’ pretraining on diverse prokaryotic genomes, which enables organism-level recognition from partial sequences.

### Meta-Embedding Construction

#### Embeddings Compression

The next part of the *non-trainable backbone* compresses the concatenation of the embeddings obtained in the previous step, reducing significantly the total number of dimensions without reducing the performance of the model. Principal Component Analysis (PCA) is used, taking only the first 500 components as the final meta-embedding, which captures more than 99.99% of the total variance.

#### Noisy Embeddings

Finally, to improve the generalization capabilities of FoundedPBI and reduce the risk of overfitting, we add a random gaussian noise to the meta-embeddings during training, with a standard deviation of 0.05. This follows the NEFTune strategy (Jain et al., 2023), and we observed an improvement in the robustness of the system when using it.

### Trainable Classification Head

The FoundedPBI framework allows us to use any model as a classification head. For the model used in the CI4CB data, a Multi-Layer Perceptron (MLP) is used, with two hidden layers of 256 and 128 neurons, and dropout layers between all of them with a dropout rate of 0.4, as it obtained the best results during the development.

This is the only trainable component of the entire system, and we trained it for 100 epochs, with a learning rate of 0.001 and a weight decay of 0.0001, using 10-Fold Cross Validation on the *train* dataset.

### Evaluation Metrics

To evaluate the performance of FoundedPBI, we used standard binary classification metrics: Precision, Recall, and F1-Score, reported in the main text, and Accuracy, Specificity, and Matthews’ Correlation Coefficient (MCC), reported along with their formulas in Supplementary.

## Results

The FoundedPBI model is compared against our previous models, the PERPHECT *predictor module* (Ataee et al., 2020), based on CNNs, and *Distilled DNABERT* (Carlos, 2023), based on a modified DNABERT (Ji et al., 2021) combined with an LSTM module.

We also tested FoundedPBI against the current state of the art. To our knowledge, the best current model at this time for solving the PBI problem is PBIP (Ma et al., 2025). They use an architecture similar to ours, but work in the protein level, using the proteins of the organisms instead of the raw DNA sequences. They use a pretrained protein representation model, UniRep (Alley et al., 2019), to encode the sequences, and then make the prediction with a combination of a 4-layer CNN plus a GRU module and a final attention layer.

To compare it with our model, we used a dataset provided by them that they use as a benchmark, the PredPHI multi-species dataset, and we performed exactly the same training and testing splits. Additionally, we also included the results of the PredPHI model, as they were the ones that created the dataset initially.

For this benchmark, some architectural modifications were applied: the DNABERT-2 context length was reduced to 16K bp due to VRAM constraints, the classification head was replaced with a Branch MLP (two parallel branches of 256–128 neurons merged into a 256-neuron layer, dropout 0.4), and PCA compression was omitted. Validation splits were constructed at the organism level, ensuring each bacteriophage appears in only one fold.

### Ensemble Learning Outperforms Individual Models

We first validate our core hypothesis: that combining multiple foundation models trained on different genomic data yields superior performance compared to any single model.

To verify that the three foundation models capture complementary genomic features, we compared their embedding geometries on the PredPHI phage sequences. Figure 2 shows a joint UMAP projection of phage embeddings, revealing distinct structures across the three representations. Quantitatively, Spearman correlation between pairwise distance matrices confirms partial but incomplete agreement: Nucleotide Transformer v2 and DNABERT-2 are the most similar (*ρ* = 0.55), while MegaDNA is the most dissimilar from both (*ρ* = 0.44 and *ρ* = 0.39). This supports the hypothesis that models trained on different genomic data encode partially orthogonal features, providing a concrete motivation for the ensemble approach.

**Figure 2.**
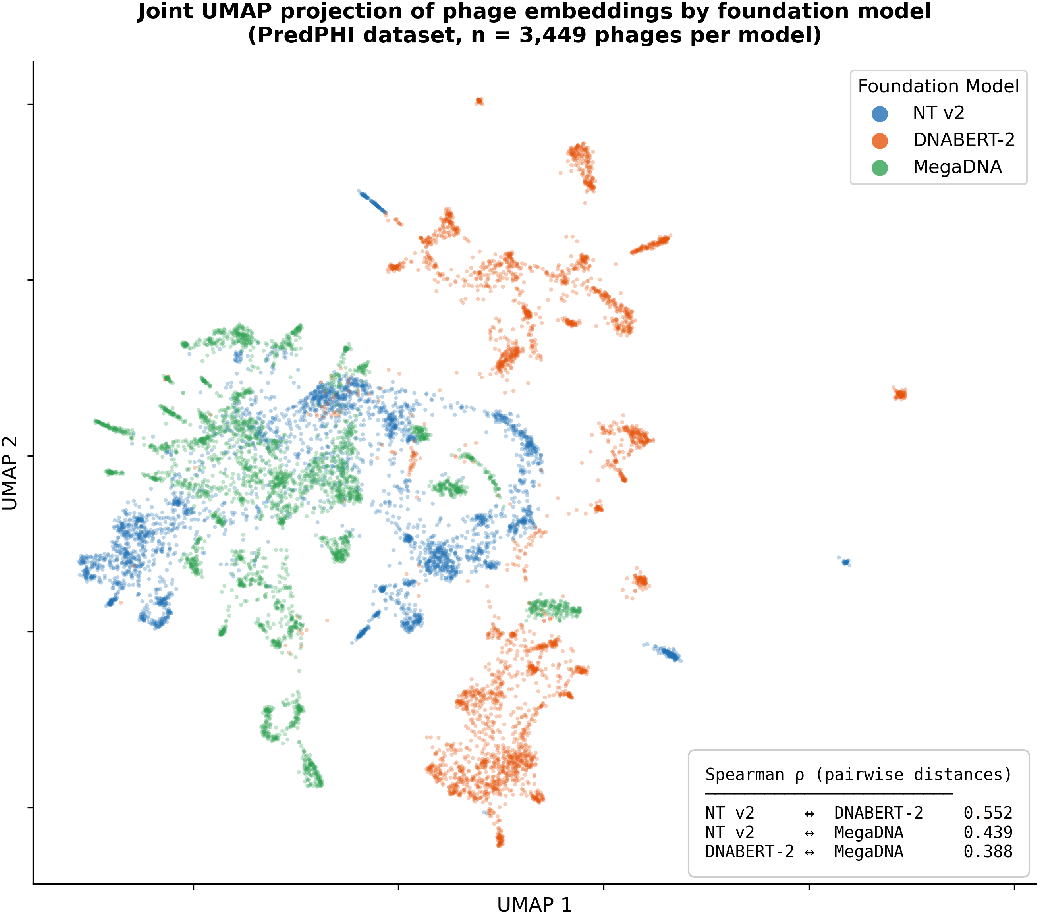
Joint UMAP projection of phage embeddings from the three foundation models. The distinct spatial organisation across models confirms that they capture complementary genomic features. Inset: Spearman *ρ* between pairwise distance matrices computed on the original embeddings.

Table 1 presents the classification results for the ensemble versus each constituent model used independently. The ensemble achieves consistent improvements over individual models on both datasets. On CI4CB, the gain is modest (+2% F1, from 0.91 to 0.93), likely due to the dataset’s limited organism diversity and smaller size, which allows even individual models to capture the dominant patterns effectively. However, on the more challenging PredPHI benchmark with 301 bacterial species and 3,449 phages, the ensemble provides substantial improvements: +6% F1 over the best individual model (MegaDNA, from 0.70 to 0.76) and +12% over the worst (DNABERT-2, from 0.64 to 0.76).

**Table 1.**
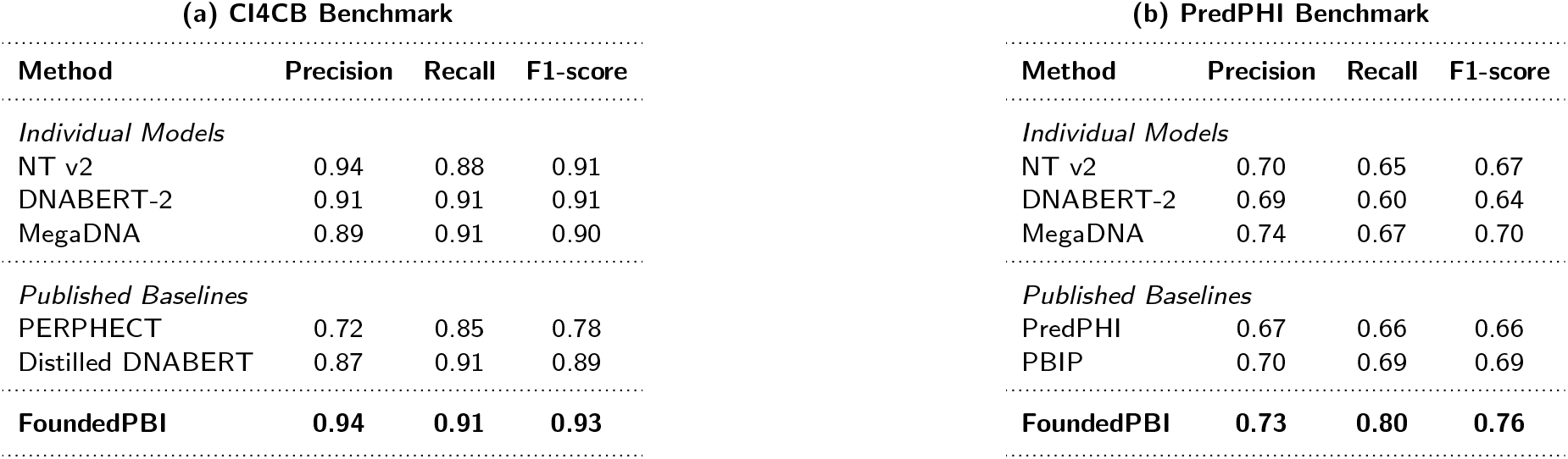
Comprehensive performance comparison on both benchmarks. FoundedPBI consistently outperforms both individual foundation models and published state-of-the-art methods. (a) CI4CB: 4% F1 improvement over our team’s previous best (Distilled DNABERT) and 2% over best individual model. (b) PredPHI: 7% F1 improvement over state-of-the-art (PBIP) and 6% over best individual model (MegaDNA).

This pattern strongly supports our hypothesis that models trained on different genomic data, prokaryotic genomes (Nucleotide Transformer v2, DNABERT-2) versus phages (MegaDNA), capture complementary features of the interaction mechanism. Interestingly, MegaDNA (trained exclusively on phages) performs best individually on PredPHI, while Nucleotide Transformer v2 and DNABERT-2 perform better on CI4CB. The ensemble successfully leverages both perspectives to achieve superior performance on both datasets.

The larger ensemble advantage on PredPHI (+6%) versus CI4CB (+2%) confirms that combining models provides greater benefits when generalizing across diverse organisms, precisely the scenario most relevant for practical phage therapy applications where novel bacteria-phage pairs must be predicted.

### Comparison with State-of-the-Art

Having established the value of ensemble learning, we compare FoundedPBI against published methods on both benchmarks presented in Table 1.

On the CI4CB dataset, FoundedPBI achieves 93% F1-score, improving over our team’s previous best method (Distilled DNABERT, 2023) by 4%. This progression—from CNN-based approaches (PERPHECT, 78% F1) to single foundation models (Distilled DNABERT, 89% F1) to ensemble foundation models (FoundedPBI, 93% F1)—demonstrates clear advancement in leveraging pre-trained genomic representations for this task.

On the PredPHI benchmark, FoundedPBI achieves 76% F1-score, outperforming the current state-of-the-art PBIP (Ma et al., 2025) by 7%, despite PBIP using task-specific protein embeddings from UniRep (Alley et al., 2019) fine-tuned on interaction data.

Notably, the improvement is particularly pronounced in recall (+11%, from 0.69 to 0.80), indicating that FoundedPBI better identifies true positive interactions. The strong performance across both benchmarks validates FoundedPBI’s generalization capabilities and practical utility for accelerating phage discovery.

### Ablation Study: Long-Context and Meta-Embedding Components

Having validated the ensemble approach, we now examine the contribution of other design choices: long-context aggregation strategies, dimensionality reduction, and embedding noise.

We adapted document-level NLP strategies to handle DNA sequences that exceed the context window of foundation models. To validate the selection of complex strategies (see Supplementary Figure S1) over naive approaches, we compared the proposed method against the simplest strategy, **Truncate Strategy**, or using only the first *n* tokens of the sequence and discarding the rest, where *n* is the maximum sequence length for each model.

Table 2 shows that using the naive truncate strategy results in a significant performance drop (−12%) in the PredPHI data, indicating that a single genomic fragment is insufficient and that combining information from multiple sequence regions improves discriminative power. As it happened with the individual foundation models, in the CI4CB data the drop is less noticeable (−2%), but the contribution can be clearly seen in the PredPHI data.

**Table 2.** Ablation study of efficacy of long-sequence strategies.

| Dataset | Model Configuration | Precision | Recall | F1-Score |
| --- | --- | --- | --- | --- |
| CI4CB | Truncate Strategy Only | 0.94 | 0.87 | 0.91 |
|  | **FoundedPBI (Best Strategy)** | **0.94** | **0.91** | **0.93** |
| PredPHI | Truncate Strategy Only | 0.69 | 0.59 | 0.63 |
|  | **FoundedPBI (Best Strategy)** | **0.73** | **0.80** | **0.76** |

Next, we analyzed the trade-off between computational efficiency and predictive performance introduced by the PCA compression step. The concatenation of raw embeddings from all three models results in a vector size of *>* 2000 dimensions, which increases training time and overfitting risk. This step is only performed in the CI4CB dataset. Applying PCA had a negligible impact on F1-Score (*<* 0.01 difference). This validates that the meta-embedding contains redundant information that can be effectively compressed without loss of relevant data.

Finally, we also explored the effect that adding noise to the embeddings had to the performance of the model. In the CI4CB dataset, similarly to before, the effect can barely be observed (*<* 0.01 difference in F1-Score). However, in the PredPHI dataset, without adding the noise, it achieves a Precision, Recall and F1-Score in test of 0.73, 0.67, 0.70, respectively, confirming that the added noise has a significant impact in the generalization capabilities and final result of FoundedPBI.

### Taxonomic Error Analysis

To better understand the limitations of FoundedPBI, we performed a taxonomic breakdown of prediction errors on the PredPHI test set (1,236 pairs, 308 errors, 24.9% overall error rate). Supplementary Table S3 reports performance grouped by bacterial host family (families with ≥20 test pairs).

Performance varies substantially across families. Streptomycetaceae achieves the lowest error rate (10.6%, *n*=142), while Pseudomonadaceae exhibits the highest (50.9%, *n*=112). Despite comparable sample sizes, these families differ nearly fivefold in error rate, suggesting that prediction difficulty is driven by biological factors rather than data availability alone. At the species level, *Pseudomonas aeruginosa* alone accounts for 53 of 308 errors (17.2%), with a false negative rate of 50.5%. The three highest-error species (*P. aeruginosa, A. baumannii, V. cholerae*) contribute 25.0% of all errors yet represent only 12.6% of test pairs—all are Gammaproteobacteria with well-documented variability in phage receptor structures (Bertozzi Silva et al., 2016).

Failure modes also differ across families: Pseudomonadaceae and Vibrionaceae errors are dominated by false negatives (51.5% and 45.2%), indicating the model under-predicts interactions, while Corynebacteriaceae and Rhizobiaceae show predominantly false positives (70.0% and 46.9%). On the phage side, nearly all test phages belong to Caudoviricetes, limiting taxonomic resolution; at the family level, Kyanoviridae (61.5%) and Autoscriptoviridae (59.5%) show the highest error rates.

## Discussion and Conclusion

We introduced FoundedPBI, a modular ensemble framework that leverages genomic foundation models and NLP-inspired long-context strategies to predict phage-bacterium interactions from DNA sequence alone. On our internal CI4CB dataset, FoundedPBI significantly outperforms previous approaches, and on the PredPHI benchmark it surpasses the current state-of-the-art.

Furthermore, while the current system utilizes complex meta-embeddings to maximize accuracy, an important objective for future development is the interpretation of these embeddings to ensure biological interpretability. We intend to implement attention-based extraction methods to map the model’s regions of interest back to the source DNA, as we anticipate that the model is implicitly learning to recognize specific interaction motifs.

Our taxonomic error analysis reveals that prediction difficulty is strongly family-dependent, with clinically critical Gram-negative families (Pseudomonadaceae, Moraxellaceae) exhibiting error rates 3–4 times higher than well-characterized Actinomycete families. This highlights a practical limitation: the pathogens most urgently targeted by phage therapy are those for which genomic sequence-based prediction remains least reliable, likely because their phage susceptibility depends on variable surface structures not fully captured by reference genome embeddings.

Finally, with this work we also showed that multiple NLP techniques can be applied to the genomics field, such as dealing with long documents or combining embeddings from multiple sources.

## Data and Code availability

The code is available at https://github.com/CI4CB-lab/FoundedPBI-code. The CI4CB public dataset (Leite et al., 2018) was gathered from GenBank and PhagesDB and is available upon request. PredPHI interactions are available at https://github.com/a1678019300/PBIP; sequences were retrieved from NCBI RefSeq via Biopython’s Entrez module.

## Author contributions statement

**P.C.B**.: Methodology, Investigation, Software, Writing. **A.B**.: Investigation, Visualization, Writing & review. **C.A.P**.: Supervision, Conceptualization, Validation, Writing & review.

## References

E. C. Alley, G. Khimulya, S. Biswas, M. AlQuraishi, and G. M. Church. Unified rational protein engineering with sequence-based deep representation learning. Nature Methods, 16(12): 1315–1322, 10 2019. doi: 10.1038/s41592-019-0598-1. URL https://www.nature.com/articles/s41592-019-0598-1.

S. Ataee, Ó. Rodríguez, X. Brochet, and C. A. Pena. Towards bacteriophage genetic edition: Deep learning prediction of phage-bacterium interactions. In 2020 IEEE International Conference on Bioinformatics and Biomedicine (BIBM), pages 2946–2948, 2020. doi: 10.1109/BIBM49941.2020.9313487.

M. Barron, PhD. Phage Therapy: past, present and future, 8 2022. URL https://asm.org/articles/2022/august/phage-therapy-past,-present-and-future.

J. Bertozzi Silva, Z. Storms, and D. Sauvageau. Host receptors for bacteriophage adsorption. FEMS Microbiol. Lett., 363(4): fnw002, Feb. 2016.

G. Brixi, M. G. Durrant, J. Ku, M. Poli, G. Brockman, D. Chang, G. A. Gonzalez, S. H. King, D. B. Li, A. T. Merchant, M. Naghipourfar, E. Nguyen, C. Ricci-Tam, D. W. Romero, G. Sun, A. Taghibakshi, A. Vorontsov, B. Yang, M. Deng, L. Gorton, N. Nguyen, N. K. Wang, E. Adams, S. A. Baccus, S. Dillmann, S. Ermon, D. Guo, R. Ilango, K. Janik, A. X. Lu, R. Mehta, M. R. Mofrad, M. Y. Ng, J. Pannu, C. Ré, J. C. Schmok, J. S. John, J. Sullivan, K. Zhu, G. Zynda, D. Balsam, P. Collison, A. B. Costa, T. Hernandez-Boussard, E. Ho, M.-Y. Liu, T. McGrath, K. Powell, D. P. Burke, H. Goodarzi, P. D. Hsu, and B. L. Hie. Genome modeling and design across all domains of life with evo 2. bioRxiv, 2025. doi: 10.1101/2025.02.18.638918. URL https://www.biorxiv.org/content/early/2025/02/21/2025.02.18.638918.

M. G. Carlos. Deep learning on genomics using NLP-oriented algorithms, 6 2023. URL https://upcommons.upc.edu/entities/publication/4306bdb5-e7ec-4217-b011-8db6ba4b9ca9/full.

H. Dalla-Torre, L. Gonzalez, J. Mendoza-Revilla, N. L. Carranza, A. H. Grzywaczewski, F. Oteri, C. Dallago, E. Trop, B. P. De Almeida, H. Sirelkhatim, G. Richard, M. Skwark, K. Beguir, M. Lopez, and T. Pierrot. Nucleotide Transformer: building and evaluating robust foundation models for human genomics. Nature Methods, 22(2):287–297, 11 2024. doi: 10.1038/s41592-024-02523-z. URL https://www.nature.com/articles/s41592-024-02523-z.

T. G. Dietterich. Ensemble methods in machine learning. In Multiple Classifier Systems, pages 1–15, Berlin, Heidelberg, 2000. Springer Berlin Heidelberg. ISBN 978-3-540-45014-6. doi: 10.1007/3-540-45014-91.

C. N. Ellington, N. Sun, N. Ho, T. Tao, S. Mahbub, D. Li, Y. Zhuang, H. Wang, L. Song, and E. P. Xing. Accurate and general dna representations emerge from genome foundation models at scale. bioRxiv, 2025. doi: 10.1101/2024.12.01.625444. URL https://www.biorxiv.org/content/early/2025/01/11/2024.12.01.625444.

N. Jain, P. yeh Chiang, Y. Wen, J. Kirchenbauer, H.-M. Chu, G. Somepalli, B. R. Bartoldson, B. Kailkhura, A. Schwarzschild, A. Saha, M. Goldblum, J. Geiping, and T. Goldstein. Neftune: Noisy embeddings improve instruction finetuning, 2023. URL https://arxiv.org/abs/2310.05914.

Y. Ji, Z. Zhou, H. Liu, and R. V. Davuluri. Dnabert: pre-trained bidirectional encoder representations from transformers model for dna-language in genome. Bioinformatics, 37(15):2112– 2120, 02 2021. doi: 10.1093/bioinformatics/btab083. URL https://doi.org/10.1093/bioinformatics/btab083.

D. M. C. Leite, X. Brochet, G. Resch, Y.-A. Que, A. Neves, and C. Peña-Reyes. Computational prediction of inter-species relationships through omics data analysis and machine learning. BMC Bioinformatics, 19(S14):420, 11 2018. doi: 10.1186/s12859-018-2388-7. URL https://pubmed.ncbi.nlm.nih.gov/30453987/.

M. Li, Y. Wang, F. Li, Y. Zhao, M. Liu, S. Zhang, Y. Bin, A. I. Smith, G. I. Webb, J. Li, J. Song, and J. Xia. A deep learning-based method for identification of bacteriophage-host interaction. IEEE/ACM Transactions on Computational Biology and Bioinformatics, 18(5):1801–1810, 2021. doi: 10.1109/TCBB.2020.3017386.

S. E. Luria and M. Delbrück. Mutations of bacteria from virus sensitivity to virus resistance. Genetics, 28(6):491–511, Nov. 1943. doi: 10.1093/genetics/28.6.491. URL https://pmc.ncbi.nlm.nih.gov/articles/PMC1209226/.

L. Ma, P. Gao, G. Liu, Y. Bai, Q. Lin, J. Li, and M. Xiao. Pbip: a deep learning framework for predicting phage–bacterium interactions at the strain level. Briefings in Bioinformatics, 26(6):bbaf656, 12 2025. ISSN 1477-4054. doi: 10.1093/bib/bbaf656. URL https://doi.org/10.1093/bib/bbaf656.

E. Nguyen, M. Poli, M. G. Durrant, B. Kang, D. Katrekar, D. B. Li, L. J. Bartie, A. W. Thomas, S. H. King, G. Brixi, J. Sullivan, M. Y. Ng, A. Lewis, A. Lou, S. Ermon, S. A. Baccus, T. Hernandez-Boussard, C. Ré, P. D. Hsu, and B. L. Hie. Sequence modeling and design from molecular to genome scale with evo. Science, 386(6723):eado9336, 2024. doi: 10.1126/science.ado9336. URL https://www.science.org/doi/abs/10.1126/science.ado9336.

N. Oyenuga, J.F. Cobo-Díaz, A. Alvarez-Ordóñez, and E.-A. Alexa. Overview of Antimicrobial Resistant ESKAPEE Pathogens in Food Sources and Their Implications from a One Health Perspective. Microorganisms, 12(10):2084, 10 2024. doi: 10.3390/microorganisms12102084. URL https://pmc.ncbi.nlm.nih.gov/articles/PMC11510272/.

D. A. Russell and G. F. Hatfull. PhagesDB: the actinobacteriophage database. Bioinformatics, 33(5): 784–786, 11 2016. doi: 10.1093/bioinformatics/btw711. URL https://pmc.ncbi.nlm.nih.gov/articles/PMC5860397/.

S. Sannigrahi, J. van Genabith, and C. Espana-Bonet. Are the best multilingual document embeddings simply based on sentence embeddings?, 2023. URL https://arxiv.org/abs/2304.14796.

E. W. Sayers, M. Cavanaugh, K. Clark, K. D. Pruitt, C. L. Schoch, S. T. Sherry, and I. Karsch-Mizrachi. Genbank. Nucleic Acids Research, 49(D1):D92–D96, 11 2020. ISSN 0305-1048. doi: 10.1093/nar/gkaa1023. URL https://doi.org/10.1093/nar/gkaa1023.

B. Shao and J. Yan. A long-context language model for deciphering and generating bacteriophage genomes. Nature Communications, 15(1):9392, 10 2024. doi: 10.1038/s41467-024-53759-4. URL https://www.nature.com/articles/s41467-024-53759-4.

L. Yu, D. Simig, C. Flaherty, A. Aghajanyan, L. Zettlemoyer, and M. Lewis. Megabyte: Predicting million-byte sequences with multiscale transformers, 2023. URL https://arxiv.org/abs/2305.07185.

F. Zhou, Y. He, J. Bai, Y. Gao, and Y. Wang. Viper: Virus-prokaryote interaction prediction using foundation model with hierarchical contrastive learning. In 2024 IEEE International Conference on Bioinformatics and Biomedicine (BIBM), pages 597–602, 2024a. doi: 10.1109/BIBM62325.2024.10822305.

Z. Zhou, Y. Ji, W. Li, P. Dutta, R. Davuluri, and H. Liu. Dnabert-2: Efficient foundation model and benchmark for multi-species genome, 2024b. URL https://arxiv.org/abs/2306.15006.

