## Supplementary files for "FoundedPBI: Using Genomic Foundation Models to predict Phage-Bacterium Interactions"

Pere Carrillo Barrera et al.

---

### S1 Supplementary Figures

Best strategies (f1 >= 0.9227: 367 exps)

| | NT2PHAGESTRAT | MEGADNAPHAGESTRAT | DNABERTPHAGESTRAT | NT2BACTSTRAT | MEGADNABACTSTRAT | DNABERTBACTSTRAT |
|---|---|---|---|---|---|---|
| BottomTruncateStrategy | 3.27% | 12.81% | 10.08% | 3.27% | 32.15% | 41.14% |
| MaxStrategy | 2.18% | 25.07% | 17.98% | 0.00% | 23.43% | 5.45% |
| TKPertStrategy | 42.23% | 23.71% | 24.52% | 0.00% | 0.82% | 0.00% |
| TopBottomTruncateStrategy | 44.41% | 16.08% | 25.61% | 96.46% | 23.16% | 44.69% |
| TruncateStrategy | 7.90% | 22.34% | 21.80% | 0.27% | 20.44% | 8.72% |

**Figure S1:** Best performing strategies. Composition of the top 5% experiments on a grid search optimizing the validation F1-Score, showing in green the strategies that appear more in the top and in red the ones that appear less, in a column wise gradient.

### S2 Evaluation Metrics

All metrics are defined in terms of the four entries of the confusion matrix: true positives (TP), true negatives (TN), false positives (FP), and false negatives (FN).

$$\text{Accuracy} = \frac{TP + TN}{TP + TN + FP + FN} \tag{S1}$$

$$\text{Precision} = \frac{TP}{TP + FP} \tag{S2}$$

$$\text{Recall} = \frac{TP}{TP + FN} \tag{S3}$$

$$\text{Specificity} = \frac{TN}{TN + FP} \tag{S4}$$

$$\text{F1-Score} = \frac{2 \cdot \text{Precision} \cdot \text{Recall}}{\text{Precision} + \text{Recall}} = \frac{2\,TP}{2\,TP + FP + FN} \tag{S5}$$

$$\text{MCC} = \frac{TP \cdot TN - FP \cdot FN}{\sqrt{(TP + FP)(TP + FN)(TN + FP)(TN + FN)}} \tag{S6}$$

### S3 Supplementary Tables

| Model | Accuracy | Precision | Recall | Specificity | F1-Score | MCC |
|---|---|---|---|---|---|---|
| PERPHECT Predictor | 0.85 | 0.72 | 0.85 | 0.84 | 0.78 | 0.67 |
| Distilled DNABERT | 0.86 | 0.87 | 0.91 | - | 0.89 | - |
| FoundedPBI | **0.93** | **0.93** | **0.91** | **0.94** | **0.93** | **0.86** |

**Table S1:** All evaluated metrics for the models of the CI4CB team in the test dataset from the CI4CB data

| Model | Accuracy | Precision | Recall | Specificity | F1-Score | MCC |
|---|---|---|---|---|---|---|
| PredPHI | 0.66 | 0.67 | 0.66 | **0.77** | 0.66 | 0.33 |
| PBIP | 0.69 | 0.70 | 0.69 | **0.77** | 0.69 | 0.39 |
| FoundedPBI | **0.75** | **0.73** | **0.80** | 0.70 | **0.76** | **0.51** |

**Table S2:** All evaluated metrics for the state-of-the-art models in the test split of the PredPHI dataset

**Table S3:** Prediction performance by bacterial host family on the PredPHI test set. Only families with $\geq$20 test pairs are shown. FN rate: false negative rate (missed true infections). FP rate: false positive rate (predicted interactions that do not occur).

| **Family** | **Species** | **Pairs** | **Error rate** | **FN rate** | **FP rate** |
|---|---|---|---|---|---|
| Pseudomonadaceae | 2 | 112 | 0.509 | 0.515 | 0.455 |
| Prochlorococcaceae | 1 | 23 | 0.435 | 0.429 | 0.500 |
| Corynebacteriaceae | 3 | 21 | 0.429 | 0.182 | 0.700 |
| Moraxellaceae | 3 | 44 | 0.386 | 0.370 | 0.412 |
| Rhizobiaceae | 5 | 39 | 0.385 | 0.000 | 0.469 |
| Mycobacteriaceae | 3 | 35 | 0.371 | 0.333 | 0.391 |
| Streptococcaceae | 3 | 25 | 0.360 | 0.000 | 0.450 |
| Vibrionaceae | 3 | 50 | 0.340 | 0.452 | 0.158 |
| Lysobacteraceae | 3 | 24 | 0.292 | 0.625 | 0.125 |
| Enterobacteriaceae | 5 | 42 | 0.262 | 0.278 | 0.250 |
| Yersiniaceae | 2 | 25 | 0.240 | 0.111 | 0.313 |
| Bacillaceae | 4 | 84 | 0.226 | 0.262 | 0.105 |
| Roseobacteraceae | 3 | 21 | 0.143 | 0.000 | 0.176 |
| Flavobacteriaceae | 4 | 41 | 0.122 | 0.097 | 0.200 |
| Streptomycetaceae | 5 | 142 | 0.106 | 0.053 | 0.213 |
| Propionibacteriaceae | 1 | 83 | 0.012 | 0.000 | 0.333 |
